# Evolution of spite versus evolution of altruism through a disbandment mechanism

**DOI:** 10.1101/2023.07.18.549605

**Authors:** Shun Kurokawa

## Abstract

Altruism and spite are costly to the actor; therefore, from the perspective of natural selection, the existence of them is a mysterious phenomenon and require explanation. If altruistic individuals are more likely to be recipients for altruism than non-altruistic individuals, then altruism can be favored by natural selection. Similarly, if spiteful individuals are less likely to be recipients for spite than non-spiteful individuals, then spite can be favored by natural selection. Spite is altruism’s evil twin or ugly sister of altruism. In some mechanisms (e.g. repeated interaction), if altruism is favored by natural selection, then spite is also favored by natural selection. In contrast, there is a situation which facilitates the evolution of altruism but is unlikely to facilitate the evolution of spite. Population structure is one typical example. Here we focus on the mechanism in which individuals choose to keep the interaction or stop the interaction according to the opponent’s behavior. We, by using evolutionary game theory, investigate the evolution of altruism and the evolution of spite under this mechanism. Our model shows that the evolution of spite is less likely than the evolution of altruism. This main result provides a nice intuition for why altruism is more abundant than spite in general in the real world.

## 1. Introduction

Social behavior can be divided into the four; altruism, spite, mutualism, and selfish (Hamilton, 1964, 1970). The former two are costly to the actor; therefore, through evolutionary lens, the existence of them is mysterious. Thus, the existence of altruism and the existence of spite require explanation.

If altruistic individuals are more likely to be recipients for altruism than non-altruistic individuals, then altruism can be adaptive and can be favored by natural selection (Eshel & Cavalli-Sforza, 1982; Skyrms, 1996; Fletcher & Doebeli, 2009). In a similar vein, if spiteful individuals are less likely to be recipients for spite than non-spiteful individuals, then spite can be adaptive and can be favored by natural selection (Foster et al., 2001; Gardner & West, 2004, 2006; Hamilton, 1970; Smead & Forber, 2013; West & Gardner, 2010).

Following Vickery (2003), spite is altruism’s evil twin. Following Gardner & West (2004), spite is ugly sister of altruism. Spite and altruism evolve by similar mechanisms. We will list up such three concrete mechanisms in the following. Firstly, humans or other animals interact repeatedly with the same individual. If altruists behave conditionally (i.e., do altruistic behavior towards altruists and do not do altruistic behavior towards defectors), then altruists can get benefit which defectors cannot get; therefore, altruistic behavior can be adaptive. This is called direct reciprocity (Trivers, 1971). When individuals interact repeatedly, not only altruism but also spite can evolve. This is because if spiteful individuals behave conditionally (i.e., do spiteful behavior towards non-spiteful individuals and do not do spiteful behavior towards spiteful individuals), then spiteful individuals are less harmed than non-spiteful individuals and spiteful behavior can be adaptive (Vickery, 2003). The second is reputation. If altruists get good reputation and those who have good reputation are likely to be helped, altruists can get benefit which defectors cannot get; therefore, altruism can be favored by natural selection. This is called indirect reciprocity (Nowak & Sigmund, 1998). Such usage of reputation facilitates not only the evolution of altruism but also the evolution of spite. If spiteful individuals gain reputation and people with the reputation are less likely harmed than people without the reputation, spiteful behavior can be adaptive (Johnstone & Bshary, 2004). The third is green beard effect. If altruists have green beard and altruistic individuals do altruistic behaviors selectively towards those who have green beard, then altruists are more likely to be recipients for altruism than non-altruists; therefore, altruistic behavior can be adaptive (Dawkins, 1976; Gardner & West, 2010; West & Gardner, 2010). Such green beard facilitates not only the evolution of altruism but also the evolution of spite. If spiteful individuals have green beard and spiteful individuals do spiteful behaviors selectively towards those who do not have green beard, then spiteful individuals are less likely to be recipients for spite than non-spiteful individuals; therefore, spiteful behavior can be adaptive (Bruner & Smead, 2022; Gardner & West, 2010; Madgwick, 2020; West & Gardner, 2010). As seen thus far, in the above three mechanisms, if altruism is favored by natural selection, then spite is also favored by natural selection.

On the other hand, there is a situation which facilitates the evolution of altruism but is unlikely to facilitate the evolution of spite. Population structure is one typical example. It is difficult to obtain situations in which populations are structured so that the evolution of spite is favored by natural selection (see Smead & Forber (2013)) while it is easy to obtain situations in which populations are structured so that the evolution of altruism is favored by natural selection. For example, in Kurokawa & Ihara (2013), while the condition under which altruism is favored by natural selection can be loose, the condition under which spite is favored by natural selection is very strict. Indeed, spite is less observed than altruism in general. This asymmetry may be caused by that population structure can facilitate the evolution of altruism but is unlikely to facilitate the evolution of spite. However, population structure is not necessarily unique situation which can facilitate the evolution of altruism but is unlikely to facilitate the evolution of spite. Actually, it is possible that another situation may have caused the phenomenon that spite is rare in general.

This paper focuses on the mechanism in which individuals choose to keep the interaction or stop the interaction according to the opponent’s behavior (Baumard et al., 2013; Fulker et al., 2021; Křivan & Cressman, 2020; Mathew & Boyd, 2009; Zheng et al., 2017). Zheng et al. (2017) has revealed that this mechanism facilitates the evolution of altruism. In this paper, we, by using evolutionary game theory, examine whether the mechanism facilitates the evolution of spite and if so, examine which the mechanism facilitates more, the evolution of altruism or the evolution of spite? We will investigate this topic.

The rest of this paper is structured as follows. In Section 2, we propose a model for social behavior with disbandment rules. In Section 3, we investigate the condition for the evolution of spite and the evolution of altruism. We also we make comparison between the likelihood of the evolution of spite and the likelihood of the evolution of altruism. Section 4 is devoted to discussion, and we make discussion and suggest future works to be undertaken.

## 2. Model

### 2.1. Game setting

Strategy 1 makes an action and through the action, the payoff of the player who makes the action increases by *p* and the payoff of the receiver increases by *q*. Strategy 2 does nothing. Suppose two individuals interact and each has the opportunity to adopt Strategy 1 or Strategy 2. Here, we assume that payoffs are additive, *p* + *q* (or *p*) is the payoff of a player adopting Strategy 1 when he plays against a player adopting Strategy 1 (or a player adopting Strategy 2), and, similarly, *q* (or 0) is the payoff of a player adopting Strategy 2 when he plays against a player adopting Strategy 1 (or a player adopting Strategy 2). When *p, q* > 0, Strategy 1 is mutualistic. When *p* > 0 and *q* < 0, Strategy 1 is selfish. When *p* < 0 and *q* > 0, Strategy 1 is altruistic. When *p, q* < 0, Strategy 1 is spiteful. Evolutionary biologists are interested in how behavior which is costly to the actor could have evolved. Therefore, throughout this paper, we assume

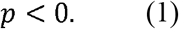

### 2.2. Strategies

We consider two strategies; one is Strategy 1 and the other is Strategy 2. Here, the population consists of individuals adopting Strategy 1 and individuals adopting Strategy 2. By *p*_*i*_, the frequency of strategy *i* is denoted. Here, we have

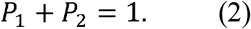

### 2.3. Pair composition

It is assumed that the population size is infinitely large. In the first round (*T* = 1), where *T* is a round number, two individuals are randomly selected from the population and matched, and two individuals in a pair play the game. By *p*_*ij*_(*T*), the frequencies of interaction pairs of strategy *i* and strategy *j* are denoted at round *T*. Note that *P*_*i*_ is independent of. *T* Let us consider a situation where Strategy 1 and Strategy 2 are present. Here, from our assumption, we have

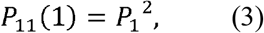

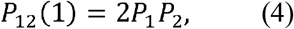

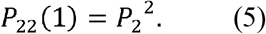

In later periods, the pair composition depends on the outcomes of the previous period. It is assumed that a probability that a pair breaks up in a round is determined by the number of Strategy 1in the dyads. The expected probability that an interaction pair *ij* will be disbanded is denoted by *ρ*_*ij*_. It is assumed that *ρ*_*ij*_ is either 1 or *ρ*, where

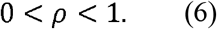

*ρ* is a background probability that a pair will naturally disband. *ρ* > 0 means that the expected number of rounds between a pair of individuals is finite. The disbandment rule can be expressed as {*ρ*_*11*_, *ρ*_*12*_, *ρ*_*22*_}. The number of disbandment rules is 8 = (2^3^). The present study investigates the dynamics under the 8 disbandment rules. The individuals who do not stop the interaction keep playing together. In contrast, those who stop interacting become single and immediately form new interaction pairs through random meetings, without paying any cost for finding a new partner.

Given that all members from disbanded pairs join the ungrouped pool, the proportion of individuals in the ungrouped pool at the end of round *T* is equal to the total proportion of pairs that were disbanded

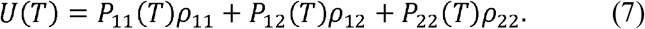

The proportion of the ungrouped pool that are cooperators is

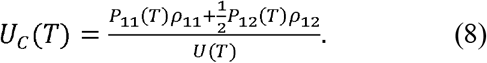

Then, for *T* ≥ 1, we can describe

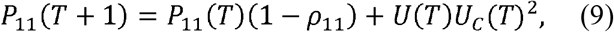

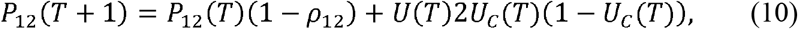

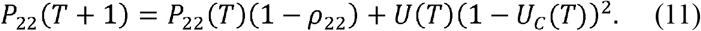

Following the above equations (3), (4), (5), (9), (10), and (11), a series of *p*_11_(*T*), *p*_12_(*T*), and *p*_22_(*T*) are determined recursively. It is assumed that such a pair formation repeats infinite times.

### 2.4. Change of strategy frequencies

φ_*j*|*i*_ (*T*) denotes the probability that a -individual has an opponent -individual at round *T*. *F*_*i*_(*T*) denotes the expected payoffs of strategy *i* at round *T*. *F*_*1*_(*T*)and *F*_*2*_(*T*) can be described as

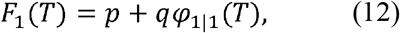

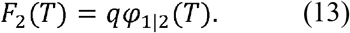

Based on previous studies (Zhang et al., 2016; Zheng et al., 2017), we assume a separation of time scales such that interaction dyads reach equilibrium quickly compared to the timescale of strategy evolution. Under this assumption, the pair formation is at an equilibrium state every time the strategy frequencies change.

It is assumed that the time evolution of *P*_*i*_ obeys a replicator equation (cf. Taylor & Jonker, 1978). Then, the replicator equation for *P*_1_ is given by

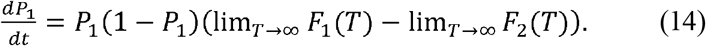

## 3. Result

### 3.1. The condition under which Strategy 1 gets a higher payoff than Strategy 2

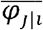 denotes the probability that a *i*-individual has an opponent *j*-individual at equilibrium, which is reached by the pair formation. Here, 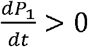 is satisfied if and only if

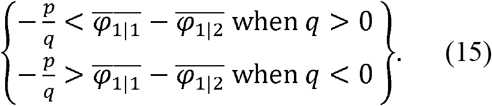

After algebraic calculation (see Appendix A for proof), we derive

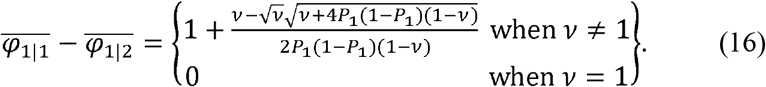

where

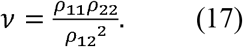

### 3.2. The exploration for 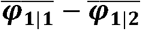

As seen in the previous section, the relationship between 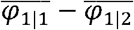 and 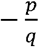 determines the dynamics. In this subsection, we will explore 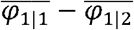. From (16), 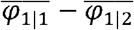 is in fluenced by *P*_1_ and *ν*. From (16), we have

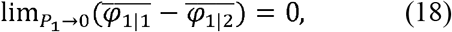

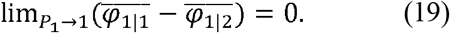

From (16), we have

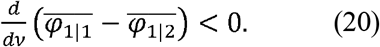

From (16), we have

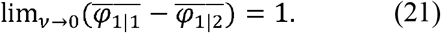

From (16), we have

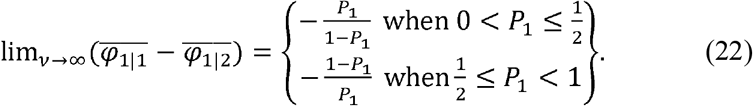

First, we investigate the case of *ν* = 1. From (16), we have

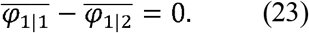

Second, we investigate the case of .*ν* < (i) 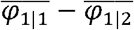 monotonically increases as *P*_1_ increases in the range of 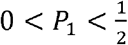, (ii) 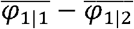 has the largest value when 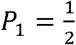 and the value of 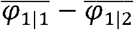 is 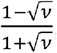, and (iii) 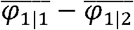 monotonically decreases as *P*_1_ increases in the range of .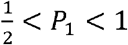. (See also Figure 1, which shows the relationship between *P*_1_ and 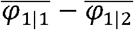).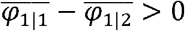 for any *P*_1_ in the range of 0. < *P*_1_ <.1. Third, we investigate the case of *ν*. >1. (i) 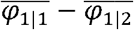 monotonically decreases as *P*_1_, increases in the range of 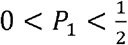, (ii) 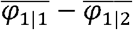 has the smallest value when 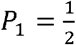 and the value of 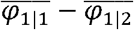 is 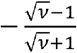 and (iii) 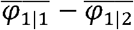 monotonically increases as *P*_1_ increases in the range of 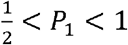 (See also Figure 1) 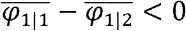 for any *P*_1_ in the range of 0. < *P*_1_ < 1

**Figure 1.**
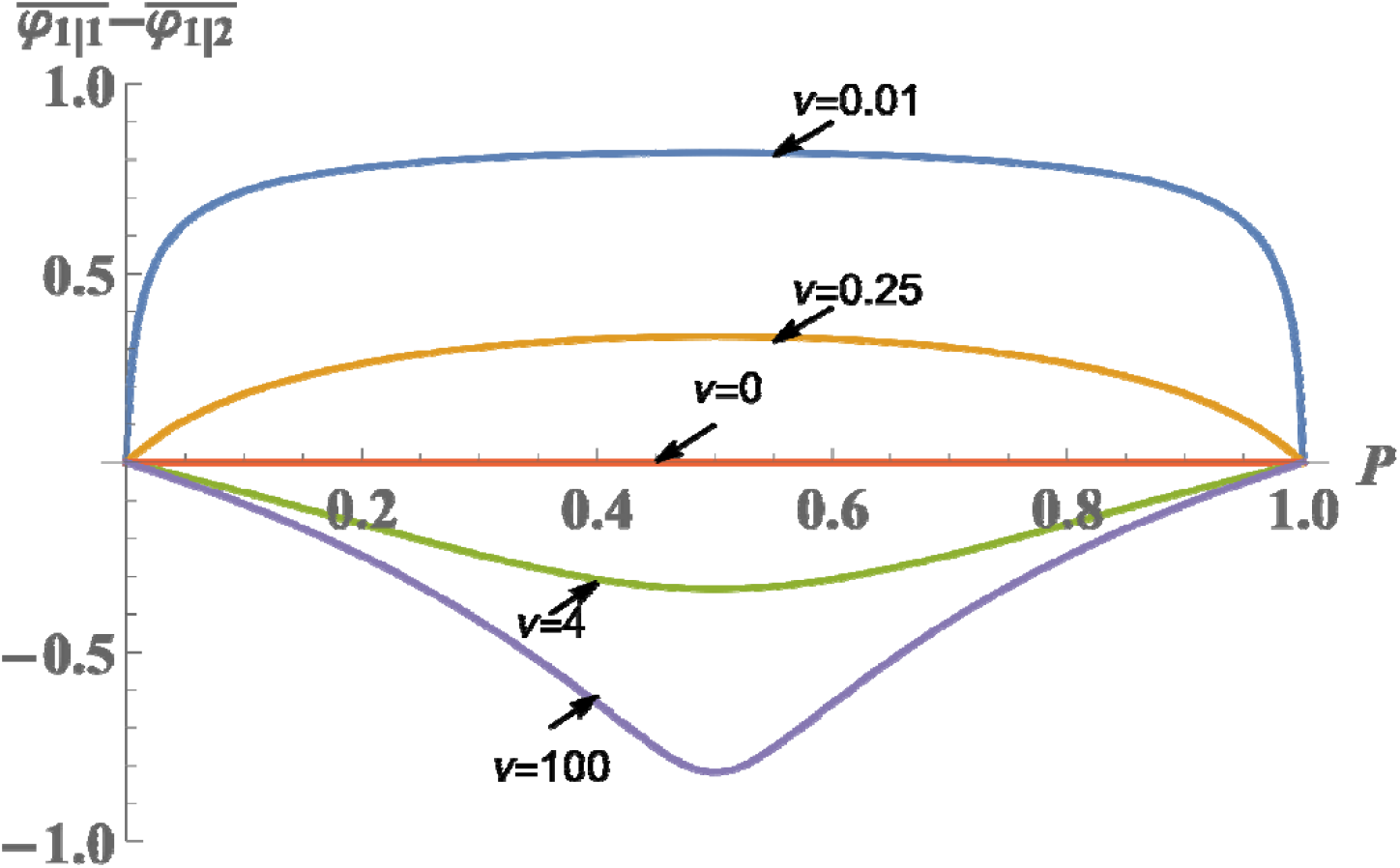
Relationship between *P*_1_ and 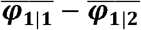. Figure 1 illustrates the relationship between *P*_1_ and 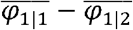, with *P*_1_ indexed on the horizontal axis and 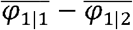 shown on the vertical axis. Figure 1 is derived from (16).

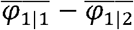 is.a function of *ν* and *P*_1_. Here, we introduce *r* (*ν,P*_1_):

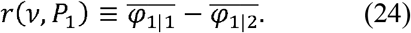

We can algebraically show that for 0. < *ν*^*^ < 1 with *ν*^*^ *ν*^*\*\**^ =1,

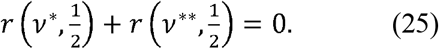

(25) is equivalent to

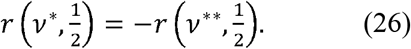

(26) means that 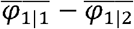 in the case where 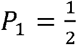 for *ν* = *ν*^*^ and 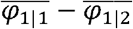 in the case where 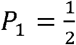 for *ν* = *ν*^*^ ^*^ have the same absolute value; however, have the opposite sign when *ν*^*^ *ν*^*\*\**^ =1. For example,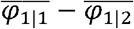 in the case where 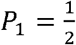 for *ν* = 0.01 and 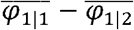 in the case where 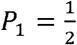 for *ν* = 100 have the same absolute value; however, have the opposite sign (see figure 1 for confirmation). For example,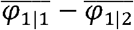 in the case where 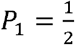 for *v =* 0.25 and 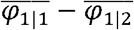 in the case where 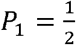 for *ν* = 4 have the same absolute value; however, have the opposite sign (see figure 1 for confirmation).

On the other hand, we can algebraically show that for *ν* ^*^ = 1 with *ν* ^*^*ν* ^*\* \**^ *=* 1 and 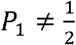, we have

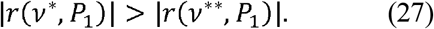

(27) means that the absolute value of 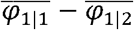 in the case where 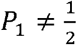 for *ν* = *ν* ^*^ is larger than the absolute value of 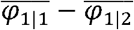 in the case where 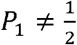 for *ν* = *ν* ^*^ when *ν* ^*^ *ν* ^* *^ *=* 1 with *ν*^*^. <1. Thus, there is asymmetry.

For example, the absolute value of 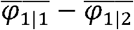 in the case where 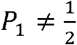 for *ν* = 0.01 is larger than the absolute value of 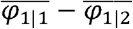 in the case where 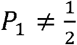 for *ν* = 100. The absolute value of 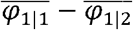 in the case where 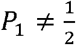 for *ν* = 0.25 is larger than the absolute value of 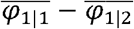 in the case where 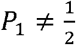 for *ν* = 4.

### 3.3. The exploration for the dynamics

In the previous subsection, we analyzed 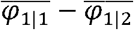. This is important, but the dynamics is determined by the relationship between 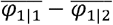 and 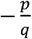. In this subsection, we will explore it.

### 3.3.1.

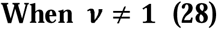

**and**

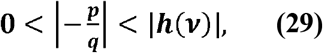

**with**

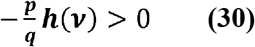

**where**

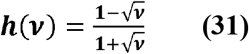

From (16), (24), and (31), we have

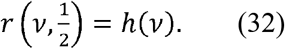

Strategy 1 gets a lower payoff than Strategy 2 when 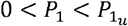 or 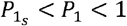. In contrast, Strategy 1 gets a higher payoff than Strategy 2 when 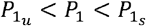, where

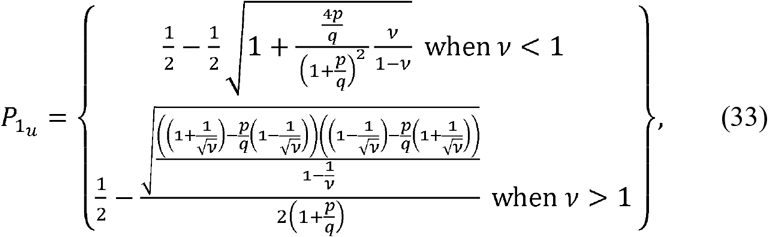

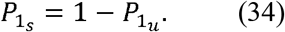

Here, 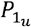 is the frequency of Strategy 1 at unstable equilibrium.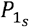 is the frequency of Strategy 1 at stable equilibrium.

When −*p/q* is near 0,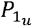 is near 0 (see figure 1). Actually, from (28) and (29), the case where −*p/q* is near 0 is classified into the case in subsection 3.3.1., as far as.*ν* ≠ 1 Besides, from (33), we algebraically derive

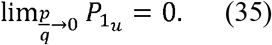

(35) shows that if *p/q* approaches 0, the basin of attraction for the population consisting of Strategy 2 shrinks to disappear no matter what is as far as.*ν* ≠ 1 Namely, when cost is small, Strategy 1 is likely to evolve. When *P*_1_ ≈ 0 is satisfied,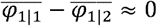. Hence, (35) can be understood naturally.

In Figure 1, we can observe that as |−*p/q*| becomes larger, the frequency of Strategy 1 at the unstable equilibrium becomes higher. Thus, the basin of attraction for the population consisting of Strategy 2 becomes larger as |−*p/q*| becomes larger. We can also observe that as |−*p/q*| becomes larger, the frequency of Strategy 1 at the stable equilibrium becomes lower.

We evaluate the likelihood of the evolution of Strategy 1 based on the following criteria: (i) The critical |−*p/q*|, above which Strategy 2 dominate Strategy 1 (i.e. Strategy 2 get higher payoffs than Strategy 1 for all *P*_1_), is high; (ii) The frequency of Strategy 1 at unstable equilibrium is low (i.e. the basin of attraction for the population consisting of Strategy 2 is small); (iii) The frequency of Strategy 1 at stable equilibrium is high (i.e. population-frequency of Strategy 1 that can persist evolutionarily is high)

No matter what is, from (31), we have

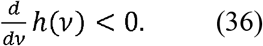

(36) holds true no matter whether *ν* is larger or smaller than 1

From (29) and (36), when *ν* < 1, as increases, the condition under which Strategy 1 is not dominated by Strategy 2 becomes strict. From (33), we have

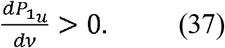

(37) means that as *ν* increases,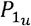 increases. By using this and (34), as increases,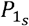 decreases. Thus, as *ν* increases, the evolution of Strategy 1 is less likely. Thus, in all the above three criteria, *ν* has a negative impact on the evolution of Strategy 1

Also, from (33), we derive

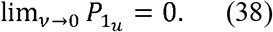

(38) shows that if approaches 0, the basin of attraction for the population consisting of Strategy 2 shrinks to disappear.

In contrast, from (29) and (36), when *v >* 1, as *ν* increases, the condition under which Strategy 1 is not dominated by Strategy 2 becomes looser. From (33), we have

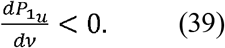

(39) means that as *ν* increases, 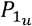 decreases. By using this and (34), as increases,*ν* increases 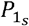. Thus, as *ν* increases, the evolution of Strategy 1 is more likely. Thus, in all the above three criteria, *ν* has a positive impact on the evolution of Strategy 1

Also, from (33), we derive

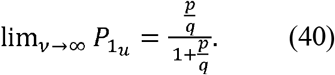

From (39) and (40), irrespective of *ν* as far as *ν* > 1, we have

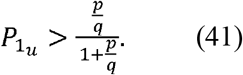

(40) and (41) show that even if *ν* increases to infinity, the basin of attraction for the population consisting of Strategy 2 does not shrink to disappear. Importantly, (38) and (40) mean that there is asymmetry between altruism and spite. We will revisit this topic later.

#### 3.3.2. Otherwise

Strategy 2 dominates Strategy 1.

### 3.4. The relationship between 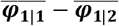 in the case of *ν* ^*∗*^ < 1 and 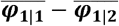 in the case of *ν* ^∗ ∗^ > 1when *ν* ^*∗*^*ν* ^*∗ ∗*^ = 1

#### 3.4.1. The condition under which Strategy 1 is not dominated by Strategy 2

From (31), we find that

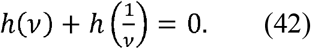

(42) shows that for *ν* ^∗^*ν* ^∗ ∗^ = 1, we have

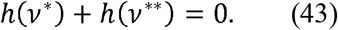

From (43), we have

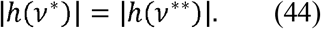

This means that under the assumption that *ν*^∗^ < 1 and *ν*^∗^ *ν*^∗ ∗^ = 1, the stringency of the condition under which altruists are not dominated by defectors in the case of *ν* = *ν*^∗^ and the stringency of the condition under which spiteful individuals are not dominated by non-spiteful individuals in the case of *ν* = *ν*^∗^are the same.

#### 3.4.2. Internal equilibria

As observed in (33), 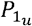 is a function of 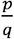 and *ν*. We have

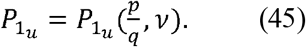

By using (33) and (45), for *t* > 0 and *ν* ^∗^ < 1, we have

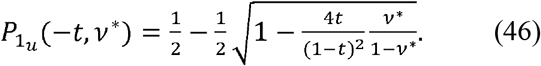

By using (33) and (45), for *t* > 0 and *ν* ^∗^ < 1, we have

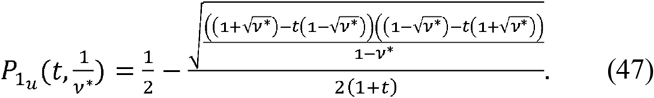

From (46) and (47), we have

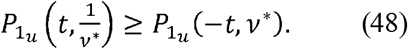

The equality in (48) holds true when and only when 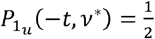. (48) shows that for, *t* > 0, *ν* ^∗^ < 1 and *ν* ^∗^*ν* ^∗ ∗^ = 1, we have

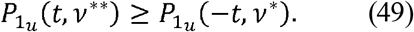

Importantly, (49) mean that there is asymmetry between the case of altruism and spite, with the basin of attraction for altruism being the greater.

### 3.5. The evolution of social behavior for each disbandment rule

In the previous section, we examine the evolution of social behavior when *ν* is given. As shown in (17), we have the relationship between *ν* and disbandment rule. Hence, we can examine the evolution of social behavior for each disbandment rule.

From (17), if the disbandment rule is and. Hence, the disbandment rule { *ρ*_11_, *ρ*_12_, *ρ*_22_} = { *ρ*,1, *ρ*}, *ν* = *ρ*^*2*^ the disbandment rule has smallest *ν* among the eight disbandment rules, irrespective of the value of *ρ* Hence, the rule { *ρ*_11_, *ρ*_12_, *ρ*_22_} = { *ρ*,1, *ρ*} most facilitates the evolution of altruism. This result is very intuitive, because altruism is facilitated when homogeneous dyads are often formed and heterogeneous dyads are rarely formed. Especially, the disbandment rule has characteristics *ρ*_11_ = *ρ* Disbandment rules that promote positive assortment promote altruism by allowing cluster of mutual cooperators to form. This result has already derived in Křivan & Cressman (2020). From (17), if the disbandment rule is { *ρ*_11_, *ρ*_12_, *ρ*_22_} = {1,1, *ρ*}, {*ρ*,1,1} *ν* = *ρ*, From (17), if the disbandment rule is disbandment rule {*ρ*_11_, *ρ*_12_, *ρ*_22_} = {1,1,1}, {*ρ,ρ,ρ*}. *ν* = *1* From (17), if the disbandment rule is {*ρ*_11_, *ρ*_12_, *ρ*_22_} = {1,*ρ,ρ*}, {*ρ,ρ*,1}, 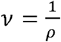. From (17), if the disbandment rule is {*ρ*_11_, *ρ*_12_, *ρ*_22_} = {1, *ρ, ρ*}, {1, *ρ*, 1}, 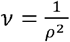. and the disbandment rule has largest *ν* among the eight disbandment rules, irrespective of the value of *ρ*. Hence, the disbandment rule {*ρ*_11_, *ρ*_12_, *ρ*_22_} = {1, *ρ*, 1} most facilitates the evolution of spite. This result is very intuitive, because spite is facilitated when homogeneous dyads are rarely formed and heterogeneous dyads are often formed. Especially, the disbandment rule has characteristics *ρ*_11_ = 1. Disbandment rules that promote negative assortment promote spite, by allowing avoiding cluster of mutually spiteful players.

We do not derive the condition under which social behavior evolves or the positions of internal equilibria for each disbandment rule. However, if we substitute the value of *ν* for each disbandment rule into (29), (33), and (34), we can obtain the condition under which social behavior evolves and the positions of internal equilibria.

Figure 2 illustrates the relationship between *P*_1_ and 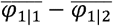 with *P*_1_ indexed on the horizontal axis and 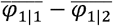 shown on the vertical axis for eight disbandment rules. (16) means that as far as *ν* is unchanged, even if {*ρ*_11_, *ρ*_12_, *ρ*_22_} is changed, 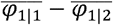 is unchanged. Actually, the lines representing the value of 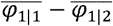 corresponding to two disbandment rules are identical if disbandment rules have the same *ν* (see figure 2).

**Figure 2.**
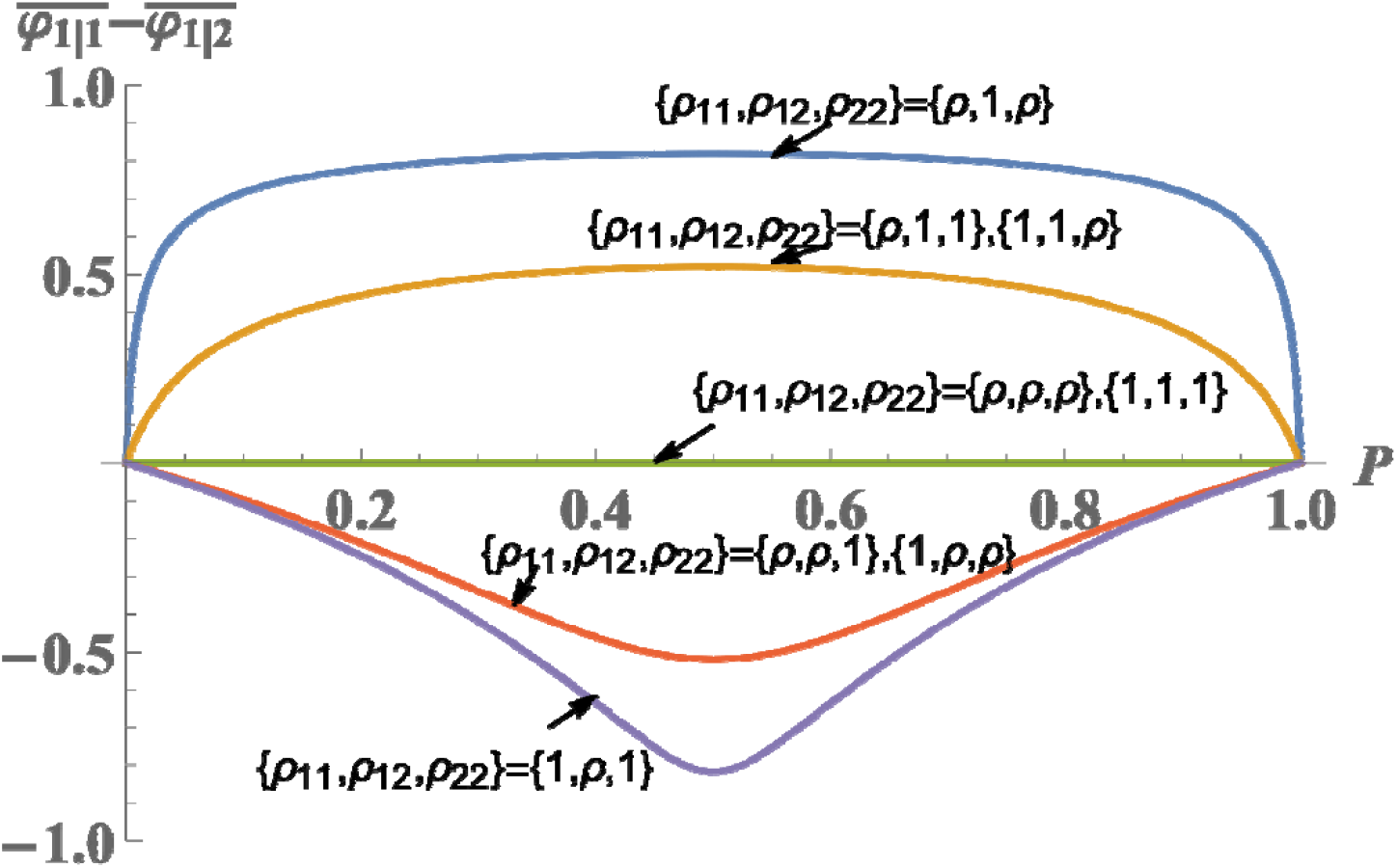
Relationship between *P*_1_ and 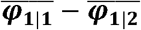 for eight disbandment rules. Figure 2 illustrates the relationship between *P*_1_ and 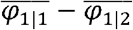, with *P*_1_ indexed on the horizontal axis and 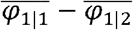 shown on the vertical axis. Figure 2 is derived from (16) and (17). The parameter value used *ρ* = 0.1.

### 3.6.The comparison between the evolution of spite and the evolution of altruism

### 3.6.1. Comparison 1

In this subsection, we make comparison between the evolution of spite in the disbandment rule which most encourages the evolution of spite and the evolution of altruism in the disbandment rule which most encourages the evolution of altruism. This comparison is on a fair basis and reasonable because we compare the best in spite with the best in altruism. When the disbandment rule is {*ρ*_11_, *ρ*_12_, *ρ*_22_} = {*ρ*,1, *ρ*}, we have

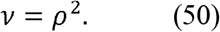

When the disbandment rule is {*ρ*_11_, *ρ*_12_, *ρ*_22_} = {1, *ρ*,1}, we have

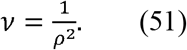

From (50) and (51), by using (44), we find that the stringency of the condition under which spiteful individuals are not dominated by non-spiteful individuals and the stringency of the condition under which altruists are not dominated by defectors are the same. On the other hand, from (50) and (51), by using (49), we find that the frequency of altruists at unstable equilibrium is lower than that of spiteful individuals at unstable equilibrium, which means that the initial evolution of altruism is more likely than that of spite. In addition, the frequency of altruists at stable equilibrium is higher than that of spiteful individuals at stable equilibrium.

#### 3.6.2. Comparison 2

In the previous subsection, we did not consider whether such disbandment rule is realized or not. However, we have no good reasons to believe that {*ρ*_11_, *ρ*_12_, *ρ*_22_} = {1, *ρ*,1} should be realized because one can easily expect that non-spiteful individuals would like to keep the pair because he/she does not receive a harm. In a similar vein, we have no good reasons to believe that {*ρ*_11_, *ρ*_12_, *ρ*_22_} = {*ρ*,1, *ρ*} should be realized because one can easily expect that defectors would like to leave the pair because he/she does not receive a benefit. In this subsection, we will care whether it can be realized or not.

Altruists get a higher payoff when interacting with altruists than when interacting with defectors. Therefore, it is natural to assume that a pair of altruists is not disbanded. Namely, *ρ*_11_ = *ρ* Defectors get a higher payoff when interacting with altruists than when interacting with defectors. Therefore, it is natural to assume that a pair of defectors is disbanded. Namely, *ρ*_22_ = 1. *ν* is minimized when *ρ*_12_ = 1 and in this case, we have

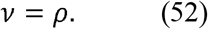

Spiteful individuals get a higher payoff when interacting with non-spiteful individuals than when interacting with spiteful individuals. Therefore, it is natural to assume that a pair of spiteful individuals is disbanded. Namely, *ρ*_11_ = 1 Non-spiteful individuals get a higher payoff when interacting with non-spiteful individuals than when interacting with spiteful individuals. Therefore, it is natural to assume that a pair of non-spiteful individuals is not disbanded. Namely, *ρ*_22_ = 1. *ν* is maximized when *ρ*_12_ = *ρ* and in this case, we have

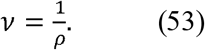

From (52) and (53), by using (44), we find that the stringency of the condition under which spiteful individuals are not dominated by non-spiteful individuals and the stringency of the condition under which altruists are not dominated by defectors are the same. On the other hand, from (52) and (53), by using (49), we find that the frequency of altruists at unstable equilibrium is lower than that of spiteful individuals at unstable equilibrium, which means that the initial evolution of altruism is more likely than that of spite. In addition, the frequency of altruists at stable equilibrium is higher than that of spiteful individuals at stable equilibrium.

## 4. Discussion

The present study examined a various disbandment rules and which disbandment rules facilitate the evolution of spite. The result was that the evolution of spite is most facilitated under the disbandment rule, where the pair of two spiteful individuals and the pair of two non-spiteful individuals are disbanded, while the pair of one spiteful individual and one non-spiteful individual is not disbanded. This result can be understood intuitively.

The evolution of spite is facilitated by disbandment rules that favor spiteful individuals and disfavor non-spiteful individuals. A rule which is good for the evolution of spite will disband groups with a composition that produces the worst payoff for spiteful individuals and the best for non-spiteful individuals, and keep groups together that produce the best payoff for spiteful individuals and the worst for non-spiteful individuals. A verbal argument based on these principles is sufficient to explain why the disbandment rule — where homogeneous dyads are disbanded and heterogeneous dyads are not — facilitates the evolution of spite for groups of size 2. Thus, although we made analysis in this paper, the result that spite is most facilitated in the disbandment rule is quite natural. In contrast, for groups of size 3 or more (note that some species interact in the groups involving more than two individuals (Joshi, 1987; Boyd & Richerson, 1988)), a verbal argument is not sufficient discern the best disbandment rule. Even for groups of size 3 or more, does the result that the evolution of spite is most encouraged when homogeneous groups are disbanded and heterogeneous groups are not still hold true? Further study on this issue may be interesting.

In addition, we investigated which is more likely to evolve, altruism or spite. The stringency of the condition under which spiteful individuals are not dominated by non-spiteful individuals and the stringency of the condition under which altruists are not dominated by defectors are the same. On the other hand, the frequency of spiteful individuals at the unstable equilibrium is higher than the frequency of altruists at the unstable equilibrium. This means that the basin of attraction for the population at which spiteful individuals and non-spiteful individuals coexist is smaller than the basin of attraction for the population at which altruists and defectors coexist. Overall, our model shows that the evolution of spite is less likely than the evolution of altruism, which is the main result in the current paper. Here, we observe that spite is rare while altruism is abundant in general in the real world (FitzGerald, 1992; Gadagkar, 1993; Pierotti, 1980; West & Gardner, 2010). This asymmetry may be caused by that population structure can facilitate the evolution of altruism but is unlikely to facilitate the evolution of spite; however, may be caused by that the mechanism in which individuals choose to keep the interaction or stop the interaction according to the opponent’s behavior can facilitate the evolution of altruism more than the evolution of spite.

The present study revealed that the evolution of spite is less likely than that of altruism with disbandment rules. How will be the result in other settings? One setting is the repeated interaction. Vickery (2003) investigated the repeated interaction and obtained the result that the evolution of spite (as well as the evolution of altruism) can be favored by natural selection in the repeated interaction and says that spite is altruism’s evil twin. However, Vickery (2003) did not make comparison between spite and altruism. Namely, Vickery (2003) did not investigate which is more likely to evolve, the evolution of spite or the evolution of altruism. Another setting is that players do not meet the same players repeatedly, but reputation is present. Johnstone & Bshary (2004) investigated the case where individuals are less likely to be harmed in the future when individuals harm others than otherwise through reputation (i.e., the spite version of indirect reciprocity). They obtained the result that the evolution of spite (as well as the evolution of altruism) can be favored by natural selection. However, Johnstone & Bshary (2004) did not make comparison between the likelihood of the evolution of spite and the likelihood of the evolution of altruism. Further study on the comparison between spite and altruism in other settings than the model setting in the current paper is warranted.

The responder’s rejecting a proposer’s offer in the ultimatum game (Harsanyi, 1961) can be regarded spite unless the amount of money is zero. It is reported that humans tend to reject such an offer that the responder can get less than half. Such a behavior has been interpreted as the reflection of human nature in which humans like fairness (i.e., humans do not want to accept a worse situation than others’). However, it may be that such a behavior should be interpreted also as the reflection of human nature in which humans are positive towards being spiteful when the cost of harm is smaller than the damage to the recipient. Actually, it can be shown that 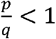 (i.e., the cost of harm is smaller than the damage to the recipient) is a condition necessary for the evolution of spite. Thus, there are (at least) two interpretations. As far as an “ordinally” ultimatum game is conducted, we are unable to reveal which interpretation is more appropriate because either type of players makes the same decision. In order to reveal which interpretation is more appropriate, we would like to propose the following modified ultimatum game experiment. One player, the proposer, is endowed with 10 dollars. The proposer kindly decides to propose that 4 dollars go to the proposer and 6 dollars go to the responder. The responder has two choices. One is accept as the proposer suggests. The other is that the responder gets 5 dollars and the proposer gets nothing. If the responder does not hope a worse situation than others’, then the responder will prefer the option in which the responder gets 6 dollars and the propose gets 4 dollars. If the responder is spiteful when and only when the cost of harm is smaller than the damage to the recipient, then the responder may prefer the option in which the responder gets 5 dollars and the propose gets nothing (note that 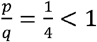 in this case). Conducting this kind of experiment will let us know that the reason why humans tend to reject such an offer that the responder can get less than half is that humans like fairness or humans are positive towards being spiteful when the cost of harm is smaller than the damage to the recipient.

## Acknowledgements

This work is partially funded by Foundation for Japan Advanced Institute of Science and Technology, Hokuriku (the JAIST Research Grants FY 2022).

## Appendix A

**Proof for (16) and (17)**

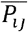 denotes the frequencies of pair *ij* at an equilibrium. We introduce

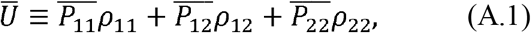

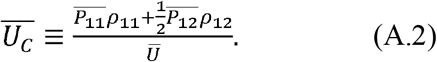

By using (9), we have

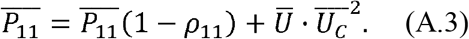

By using (10), we have

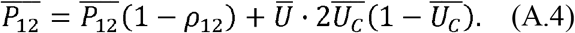

By using (11), we have

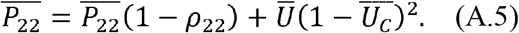

By using (A.3), (A.4), and (A.5), we derive the following relationship:

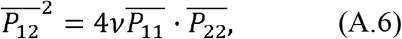

Where

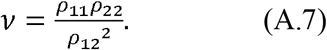

We have

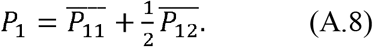

From (A.8), we have

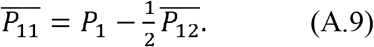

We have

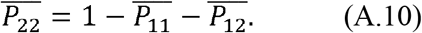

Substitution of (A.9) into (A.10) yields

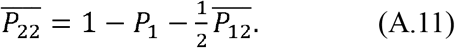

Substitution of (A.9) and (A.11) into (A.6) yields

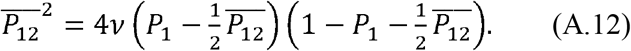

By using 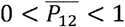 and solving (A.12) for 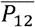, we derive

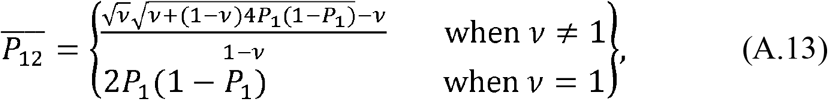

It is shown that there is just one equilibrium in the sense of pair formation.

Here, we have

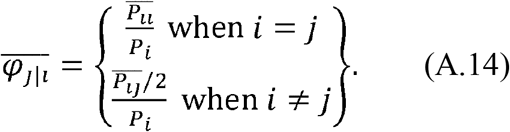

From (A.13) and (A.14), we have (16). This is the end of the proof.

